# Fitness costs of antibiotic resistance mutations in rich media do not reliably predict those in infection-like environments

**DOI:** 10.64898/2026.09.08.750013

**Authors:** Jack Knowles, Lauren M Pittaccio, Selina Lindon, Corey Woods, Danna R Gifford, Lisa J Kahl, R Craig MacLean, Simon JS Cameron, Rachel M Wheatley

## Abstract

The acquisition of antibiotic resistance is typically associated with a fitness cost, reducing the competitive ability of cells in the absence of antibiotics. Our ability to predict these costs informs our ability to predict how likely it is that resistance will subside in the absence of treatment. Fitness costs are typically measured in laboratory cultures using a nutritionally rich medium for growth. Here, we tested the impact of infection-like environments on the prevalence and magnitude of fitness costs associated with antibiotic resistance mutations in the opportunistic pathogen *Pseudomonas aeruginosa.* We assessed two different clinically relevant infection niches, the lung and urinary tract, through the use of infection-niche mimicking media that models the chemical compositions of these niches. The fitness costs of resistance mutations in rich media did not reliably predict those in infection-like environments, on average costs were increased in the lung-mimicking environment and costly mutations were less prevalent in the urinary tract-mimicking environment. Synthetic human urine significantly enhanced the fitness of *P. aeruginosa* with resistance mutations in *nfxB*, encoding a multidrug efflux pump transcriptional repressor, and we identified oxalate as the likely component driving this effect. Our findings highlight the importance of environmental chemical composition for shaping the fitness costs of resistance in *P. aeruginosa* and suggest that *nfxB* mutants may have a selective advantage in the urinary tract, mediated by the presence of oxalate.

## Introduction

Antimicrobial resistance (AMR) in pathogenic bacteria represents a critical and escalating global health threat, causing increased treatment failure and escalating healthcare costs^1^. As advances in the development and approval of new antibiotics has slowed considerably since the mid-20^th^ century^2^, it is essential to preserve the efficacy of existing treatments. Strategies like antimicrobial stewardship aim to tackle bacterial AMR by reducing unnecessary or inappropriate antibiotic use^3,4^, relying on the assumption that resistance will decline in the absence of antibiotics due to acquisition typically being associated with a fitness cost^5,6^. This fitness cost is expressed in terms of a reduced competitive ability of resistant cells compared to their antibiotic-sensitive ancestors. Longitudinal surveillance studies support that resistance can decline following reductions in antibiotic use^7,8^, but this is not always the case^9,10^. This dichotomy is also observed at the level of individual patients, where newly acquired resistance mutations can persist long after the antibiotic treatment ends^11,12^. The reasons for these divergent outcomes remain unclear.

A limitation in our current knowledge may arise from the experimental conditions under which AMR-associated fitness costs are traditionally measured. Measurements of fitness costs are conventionally conducted *in vitro* using a rich laboratory broth, such as Lysogeny Broth (LB) or Tryptic Soy Broth (TSB)^5,6^, or simple, defined media containing a single carbon source^13,14^. Whilst this provides many advantages in terms of throughput, ease, and cost, these conditions do not reflect the chemical complexities of infection niches^15,16^. *In vivo* models are an alternative approach that offer increased physiological relevance, but common animal models differ chemically from humans at key infection sites, particularly in terms of host antimicrobials ^15–18^. They also have additional drawbacks in terms of ethical considerations, higher costs, and limited scope for high-throughput investigations^19,20^.

One solution that could maintain the benefits of *in vitro* methods, whilst increasing the physiological relevance of fitness assays, is the use of infection-niche mimicking media (IMM). IMM differ from standard laboratory broths by replicating the chemical properties and compositions of infection niches, reflecting the pH, osmolarity, nutrient availability of specific infection niches and including niche-specific components like host-antimicrobials, carbon-sources, metals, and salts^15–18,21^. Environmental factors can have varied impacts on the fitness effects of antibiotic resistance mutations^13,22,23^. For example, changes in temperature have been shown to have a more profound effect on the fitness costs of rifampicin resistance mutations than changes in carbon substrate^13^.

Numerous IMM have been developed to mimic important human infection niches, such as the lung, oral cavity, urinary tract, colon, and wound sites^15,16,18,21^. As a first example, healthy lung media (HLM), developed for the study of acute lung infections, incorporates metabolites, eDNA, albumin, host antimicrobials, and mucin^16^. As a second example, synthetic human urine (SHU), developed for the study of acute urinary tract infections, reflects an acidic environment (5.6 pH), and incorporates niche-specific concentrations of salts, organic acids, nitrogenous waste, and urea^15^. Despite the increased physiological relevance of IMM, whether their use provides a tangible benefit that offsets their added cost and preparation time compared to standard laboratory broths is debated^24–29^. Previous work querying the bacterial essential genome across a variety of IMM has indicated that only a small proportion of the genome is conditionally essential for growth in IMM compared to broth (∼15-30 unique genes, ∼0.25-0.5% of genome)^24^, and gene expression can sometimes be remarkably similar between *in vitro* and *in vivo* conditions (e.g. ∼96% of genes expressed similarly in *Porphyromonas gingivalis* in the human oral cavity) ^26^.

In this work, we tested the impact of infection-like environments on the prevalence and magnitude of fitness costs associated with antibiotic resistance mutations in *Pseudomonas aeruginosa*, one of six bacterial pathogens that are collectively responsible for over 70% of the global deaths attributed to bacterial AMR^1^. We measured the fitness costs of a panel of antibiotic-resistant mutants generated spontaneously via selection on six different antipseudomonal drugs. We investigated how well fitness costs measured in TSB, a commonly used rich laboratory broth, predict those measured in two different infection-like environments, representing the lung (heathy lung media; HLM) and urinary tract (synthetic human urine; SHU) environments relevant to acute infections. These IMM represent two distinct clinically relevant infection niches for *P. aeruginosa*, which is a leading cause of ventilator-associated lung infections and indwelling catheter-associated urinary tract infections^30,31^. We find that the fitness costs of resistance mutations in rich media do not reliably predict those in infection-like environments, and oxalate present in human urine may selectively enhance the fitness of multidrug efflux pump upregulation mutants. These findings are relevant to our ability to predict how resistance may emerge and spread in different environments, and highlight the importance of the chemical composition of the infection-niche for shaping the dynamics of antibiotic resistance in *P. aeruginosa*.

## Materials and methods

### Generation of antibiotic-resistant mutants in *P. aeruginosa*

Six antibiotics (aztreonam, piperacillin/tazobactam, ceftazidime, meropenem, ciprofloxacin, and amikacin) were used to generate spontaneous antibiotic-resistant mutants from *P. aeruginosa* PAO1, as previously described^32^. The *P. aeruginosa* PAO1 strain used carries a green fluorescent protein tag (PAO1-GFP)^33^, which was used to facilitate the previously described experiments in^32^. Six independent resistant mutants were selected on each antibiotic at clinical breakpoint concentrations according to the EUCAST Clinical Breakpoints (16 μg/mL aztreonam, 16 μg/mL piperacillin-tazobactam, 8 μg/mL ceftazidime, 8 μg/mL meropenem, 0.5 μg/mL ciprofloxacin, and 16 μg/mL amikacin)^34^, following the methodology previously described in^32^. In brief, single, independent colonies of PAO1 were grown in LB Miller broth overnight (37 °C, aerobic conditions, 220 rpm shaking). Overnight cultures were then plated on LB Miller agar plates supplemented with each antibiotic at its clinical breakpoint concentration. Single colonies from each plate were then inoculated into LB Miller broth supplemented with the corresponding antibiotic at 50% of its clinical breakpoint concentration, and grown overnight before being stored in glycerol at -80 °C. Meropenem resistant mutants could not be generated using the same method; to generate meropenem resistance, PAO1 was passaged over three days on meropenem plates, and transferred every 24 h to a new plate with a higher concentration (1 μg/mL to 2 μg/mL, to 4 μg/mL, to 8 μg/mL). The same procedure for stocking as above was then followed.

### Genome sequencing and variant detection of spontaneous antibiotic resistant mutants

DNA extraction, library preparation, and sequencing of all 33 resistant strains and the ancestral PAO1 strain were carried out by MicrobesNG (Birmingham, UK), following their standard protocols. Sequencing was performed using an Illumina NovaSeq6000, following the 250 bp paired-end protocol. Adapter trimming was performed using Trimmomatic v0.30 (Q15 sliding window)^35^, and the PAO1-UW reference genome (NCBI RefSeq accession NC_002516.3) was used for alignment^36^. Variants between the ancestral PAO1 strain and resistant mutants were identified using breseq v0.36.1, with only fixed (100%) variants in resistant strains relative to the ancestral PAO1 reported (Table 1).

**Table 1:**
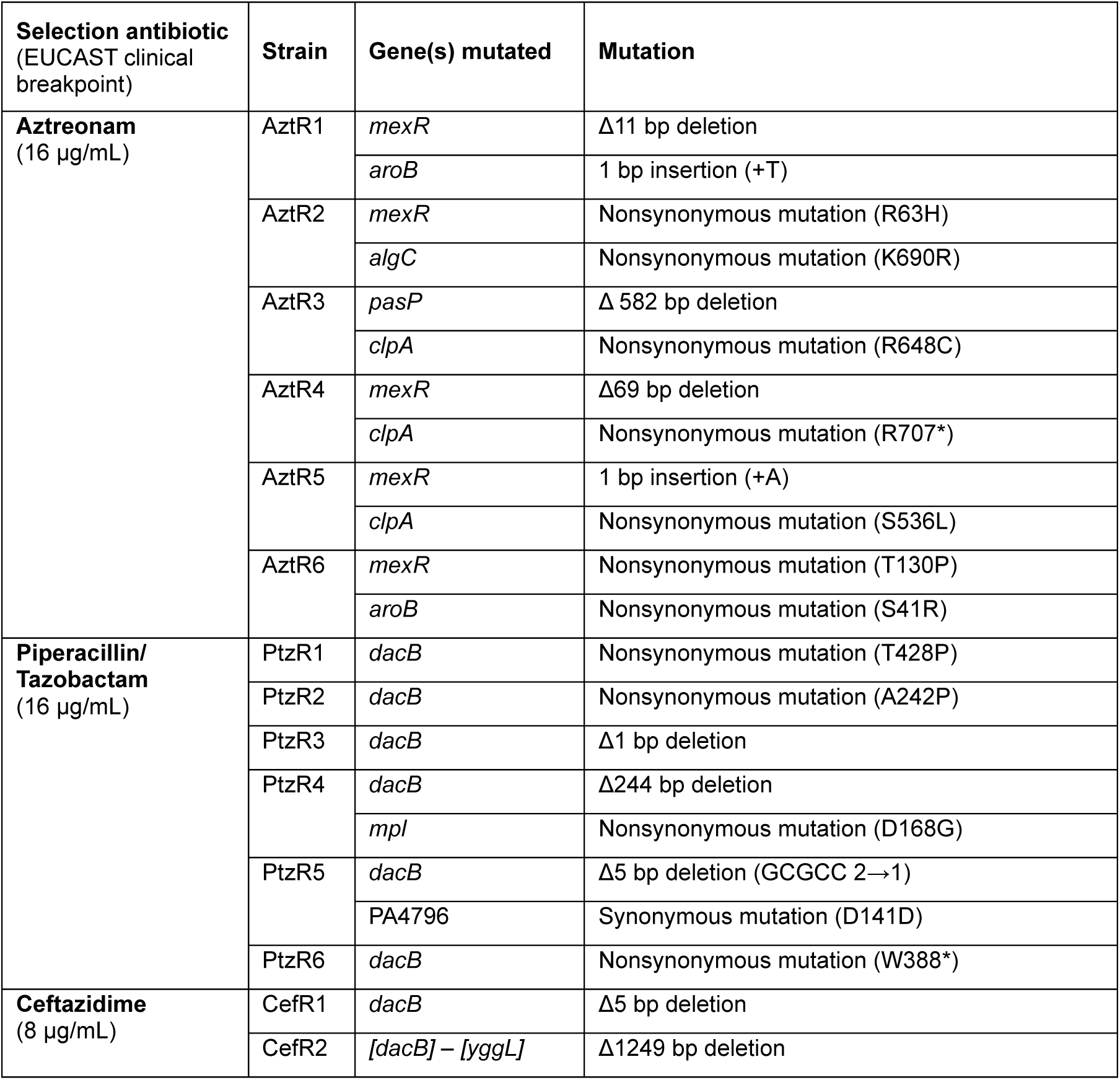
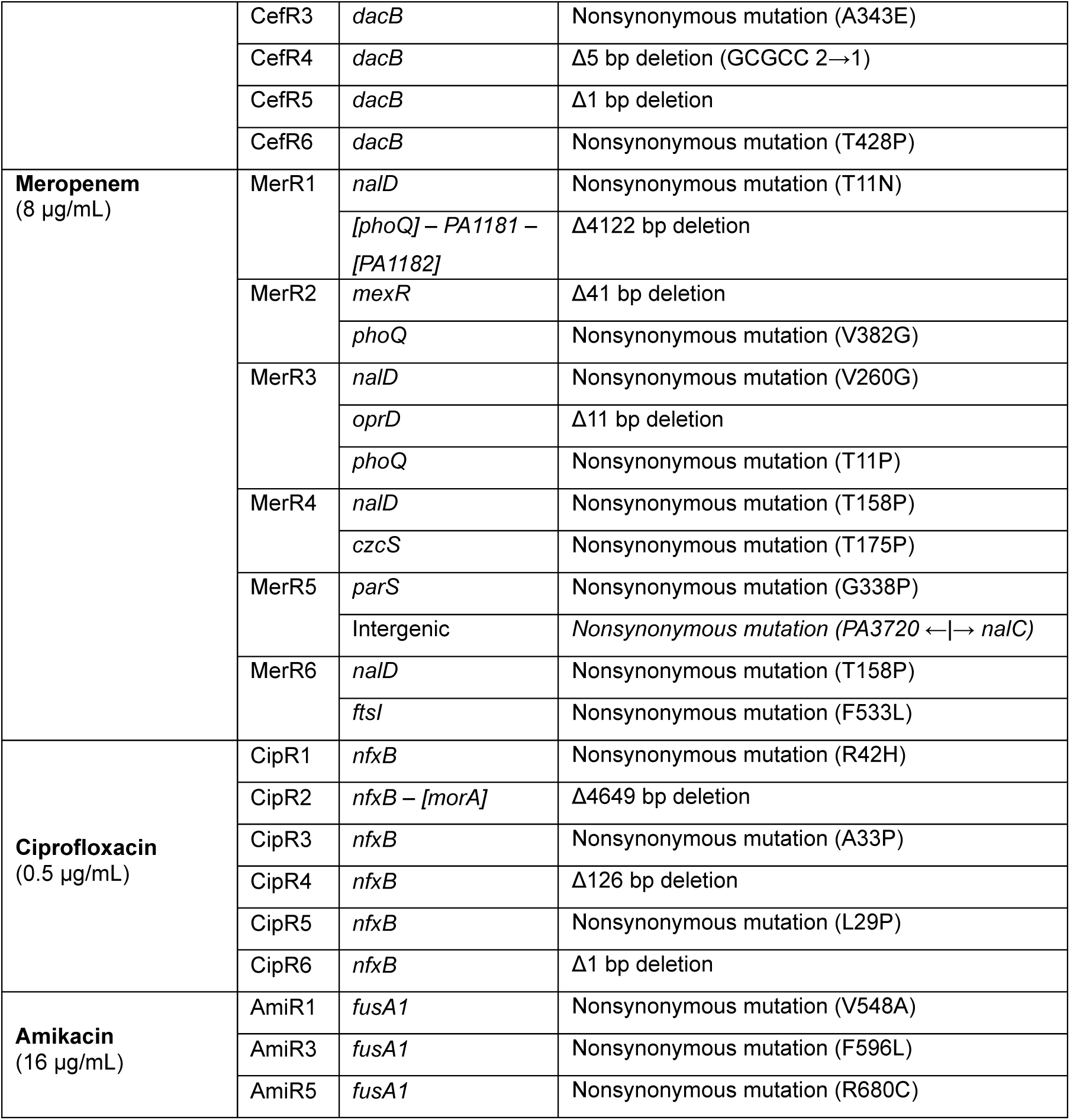
Panel of antibiotic-resistant mutants generated from the P. aeruginosa PAO1 ancestor. Description of antibiotic-resistant mutants including selection antibiotic and mutations. Square brackets indicate that a multi-gene deletion begins or ends within that gene.

### Preparation of infection-niche mimicking media

Two IMM were prepared: healthy lung media (HLM)^16^ and synthetic human urine (SHU)^15^. The compositions and pH of HLM, SHU and TSB are described in Supplementary Table S1. HLM was prepared in 2 L batches, following the protocol described in^16^, and stored in aliquots at - 80 °C until needed. SHU was prepared following the protocol described in^15^, except for the substitution of casamino acids with SC amino acid mixture (MP Biomedicals). SHU stock solutions were stored at 4 °C for up to one month, and urea and iron (II) oxide solutions were freshly prepared as required.

### Growth assays and fitness cost calculations

Strains were streaked from -80 °C glycerol stocks onto tryptic soy agar (TSA) plates supplemented with 15 μg/mL gentamicin and incubated overnight (∼18-20 h, 37 °C, static, aerobic conditions). Colonies were inoculated into the inner 60 wells of a 96-well plate containing 200 μL tryptic soy broth (TSB) and incubated overnight (∼19 h, 37 °C, 220 rpm shaking, aerobic conditions). Growth measurements were conducted in 96-well plates, with only the inner 60 wells inoculated with 5 μL of overnight culture and 195 μL of the relevant medium (TSB, HLM, or SHU). Optical density (OD_595nm_) readings were taken at 10-minute intervals for 24 hours using a FLUOStar Omega microplate reader set to 37 °C with 200 rpm double-orbital shaking for 60 seconds prior to each measurement. In each plate at least one inner well was left blank per medium to provide a blank value for growth curve analysis, and to act as a negative control. Fitness assay plates were assembled as paired comparisons split across two halves of the plate (TSB and HLM, TSB and SHU, or HLM and SHU). In each medium, six biological replicates of each mutant and 16 biological replicates of the ancestral strain were grown across all plates (plate layouts shown in Figure S1). Optical density data were analysed in RStudio, version 2024.9.0.375^37^, using the ‘growthcurver’ package^38^. The ‘SummarizeGrowth’ function was used to provide summary growth metrics per well, including area under the curve (AUC), after subtracting blank well values. Curves with a sigma value > 0.14 (top ∼5% of sigma values) were manually inspected to ensure accurate curve fitting. Out of 642 growth curves, 41 were manually inspected, and 16 were excluded due to curve shape or poor fitting of datapoints. This reduced the number of replicates for some mutants from six to five, and for the ancestral strain from 16 to 13 in TSB, and to 15 in HLM and SHU. Fitness costs as fitness relative to the ancestral strain were calculated for each mutant replicate using the following equation:

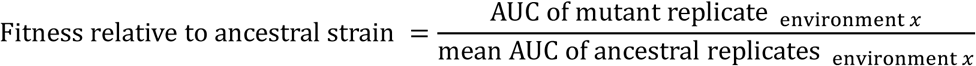

### Screening of SHU components for impact on *nfxB* mutants

To investigate the impact of SHU components on *nfxB* mutant fitness, SHU components (oxalic acid, sodium lactate, trisodium citrate, creatinine, urea, and uric acid) were dissolved in deionised, autoclaved water and filter sterilised. Oxalic acid will predominately be present in oxalate form in SHU, sodium lactate as lactate, and trisodium citrate as citrate. Individual SHU component solutions were added to TSB alongside an equal volume of 2X concentrated TSB, to produce SHU component supplemented media at the concentrations used in SHU (Table S1). pH was adjusted to 5.6 to match the pH of SHU using hydrochloric acid (0.5 M) and an equal volume of 2X TSB. SHU-component assay plates were prepared by filling the inner 60 wells of a clear 96-well plate with SHU component supplemented TSB media alongside TSB controls. Five biological replicates of three independent *nfxB* mutant strains (CipR1, CipR2, CipR4), and the ancestral PAO1 strain were prepared as overnight cultures as described above and 5 μL was transferred into each assay plate into either SHU component supplemented media or TSB, providing a within-plate volumetric control. The three *nfxB* mutants tested were chosen as they had the highest fitness in SHU (Figure 2B), and represented three different mutation types: a SNP (CipR1); a large, 4649 bp two-gene deletion of *nfxB* and *morA* (CipR2); and a smaller, 126 bp deletion that affected only *nfxB* (CipR4) (Table 1). Following growth curve analysis, AUC was used as the summary measure of growth. Fitness relative to growth in standard TSB was calculated using the following equation to investigate the impact of each SHU component on growth:

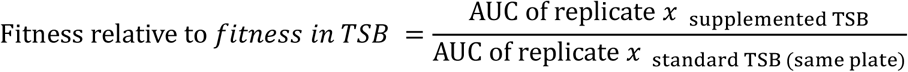

### Statistical analysis

All statistical analyses were conducted using R, version 4.4.2^39^, in RStudio, version 2024.9.0.375^37^. For correlation analyses of growth between each environment (TSB, SHU or HLM), Shapiro–Wilk tests were first used to assess the normality of AUC distributions, followed by Spearman’s rank-order correlation tests because the assumptions of Pearson’s were not met. For all subsequent analyses, ANOVA assumptions were assessed using Shapiro–Wilk tests and Q-Q plots of model residuals to evaluate normality, and Levene’s tests for assessing homogeneity of variances. For analyses comparing AUC or relative fitness across infection-like environments (TSB, HLM or SHU), Welch’s ANOVA test was conducted followed by Games-Howell post-hoc tests. For comparisons between the relative fitness of the ancestral strain and mutant strains in SHU component-supplemented media, Welch’s ANOVA was followed by Welch’s t-tests with Bonferroni corrections, used to account for multiple comparisons. All post-hoc analyses were conducted using the ‘rstatix’ package^40^.

## Results

### Characterisation of *de novo* resistance mutations in panel of *P. aeruginosa* strains

We generated 33 spontaneous resistant mutants of *P. aeruginosa* PAO1 selected at EUCAST clinical breakpoints^34^: six aztreonam resistant mutants (AztR1-R6), six piperacillin-tazobactam resistant mutants (PtzR1-ptzR6), six ceftazidime resistant mutants (cefR1-R6), six meropenem resistant mutants (MerR1-R6), six ciprofloxacin resistant mutants (cipR1-R6), and three amikacin resistant mutants (AmiR1, R3 & R5). Resistance mutations were confirmed by sequencing, by comparison to the ancestral strain (Table 1).

Each of the 33 mutants had a unique mutation or set of mutations separating them from the ancestral genotype; some mutations occurred in the same genes across strains (e.g. *dacB*, *mexR*, *nfxB*, *fusA1*, *aroB,* and *clpA*) (Table 1). The mechanisms of resistance resulting from mutations in *dacB*, *nfxB*, and *mexR* are well established in the literature: *dacB* mutations result in overexpression of the Amp-C beta-lactamase^41,42^, whereas *nfxB* and *mexR* mutations are associated with upregulation of multidrug efflux pumps^43,44^. Mutations in *fusA1,* which encodes elongation factor EF-G1A, are associated with aminoglycoside resistance^45,46^, but their effects are known to be highly pleiotropic^47,48^. Mutations in *aroB* and *clpA* were only identified in strains that also possessed a *mexR* mutation.

### Growth in TSB correlates with growth in HLM but not SHU

We conducted high-throughput growth assays of each strain in TSB, HLM, and SHU, using AUC as to summarise growth, a commonly used metric that incorporates several features of the bacterial growth curve, including growth rate and carrying capacity (Figure 1A) ^22,49,50^. As expected, the growth of the ancestral PAO1 strain (shown in red) was generally greater than that of the resistant mutants, and the majority of exceptions to this were observed in SHU (Figure 1A). To understand whether mutations impacted the growth of strains similarly in each environment, we performed correlation analyses across these three environments (Figure 1B). A significant positive correlation in growth of strains was observed between TSB and HLM (Spearman’s *rho* = 0.735, *p < 0.0001*), showing that the rank order of fitness was generally preserved between these environments. However, we detected no correlation between the growth of strains in TSB and SHU (Spearman’s rho = 0.137, *p* = 0.440), or in HLM and SHU (Spearman’s rho = 0.095, *p* = 0.591) (Figure 1B).

**Figure 1:**
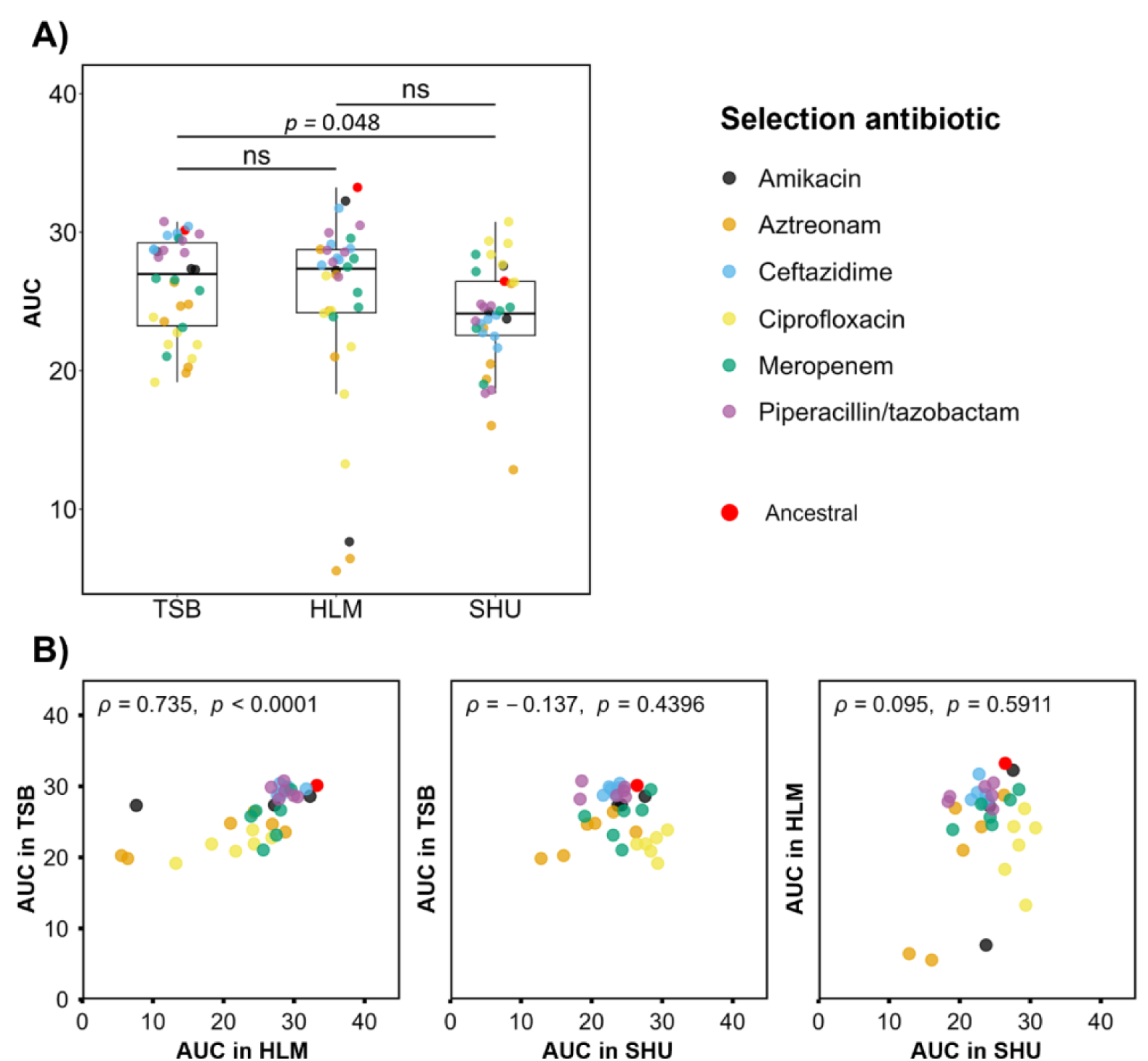
Correlation analysis of growth across TSB, HLM and SHU. **A)** Comparison of growth, shown as AUC, of all P. aeruginosa strains in each environment, Welch’s ANOVA and post-hoc Games-Howell test were used to determine significance with p-value on plot where p < 0.05. **B)** Correlation analyses compared growth, measured by AUC, of P. aeruginosa strains across all medium combinations, with Spearman’s rank correlation coefficients (rho/ρ) and p-values are shown on each plot. Each datapoint represents the mean of at least five biological replicates for every antibiotic resistant strain per environment, and at least 13 replicates for the ancestral strain.

### The prevalence and magnitude of fitness costs of resistance differ in infection-like environments

In order to determine how the fitness costs of resistance changed across environments, we compared growth (AUC) of each mutant relative to the ancestral strain in each environment (Figure 2A). Resistance mutations were, on average, more costly in HLM, and costly mutations were less prevalent in SHU (Figure 2A). When comparing the growth (AUC) of individual mutants to the ancestral strain within environments, in TSB 13/33 (∼39%) of mutants had significantly reduced fitness compared to the ancestral strain, this was 19/33 (∼58%) of mutants in HLM, and 6/33 (∼18%) of mutants in SHU (Figures 2B and S2). Notably, all ciprofloxacin resistant mutants (CipR1-R6) were found to have significant fitness costs in TSB, but none exhibited significant fitness costs in SHU and in fact for 5/6 mutants relative fitness in SHU trended higher than the ancestral strain (Figures 2B and S2). The magnitude of fitness costs varied between TSB and the infection-like environments (Figure 2B). In TSB, no mutants incurred costs that reduced growth by more than 40%, in HLM there were 5/33 (∼15%) and in SHU there were 2/33 (∼6%). Conversely more mutants had an AUC within 10% of the ancestral strains in TSB (16/33 or ∼48%) and SHU (17/33 or ∼52%) than in HLM (4/33 or ∼12%) (Figure 2B).

**Figure 2:**
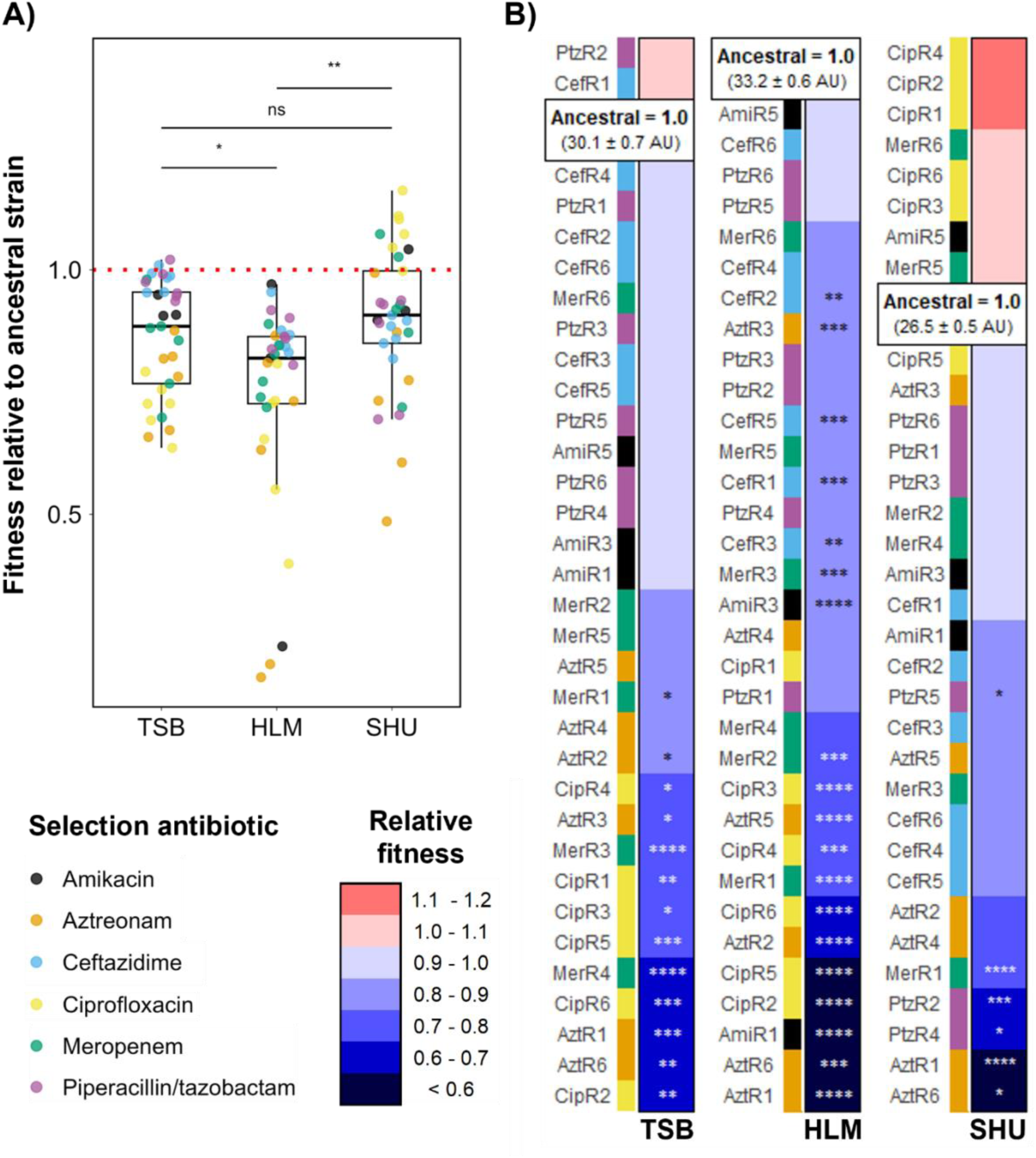
Prevalence and magnitude of fitness costs associated with antibiotic resistance mutations in P. aeruginosa across TSB, HLM, and SHU. **A)** Comparison of the fitness costs of resistance in each environment, as assessed by fitness relative to the ancestral strain (shown by red dotted line). **B)** Comparison to ancestral growth (AUC) on a per environment, per mutant basis, data visualised as fitness relative to the ancestral strain with the AUC and SEM shown for the ancestor, and with data rank ordered and colour-coded by mean relative fitness. Statistical significance was assessed in both A and B using Welch’s ANOVA followed by a Games–Howell post-hoc test. Significance annotations: p < 0.05 (*), p < 0.01 (**), p < 0.001 (***), p < 0.0001 (****).

### Synthetic human urine alleviates the fitness costs associated with *nfxB* mutations

While each resistant mutant possessed unique mutations, a subset of 17 mutants had resistance conferred by a single mutation in one of three genes (Table 1). These three genes were *dacB, nfxB*, *and fusA1*, frequent mutational targets in clinical isolates^47,51–54^. We grouped the 17 strains by mutated gene to determine whether environment alters the fitness costs associated with resistance mutations on a per gene basis (Figure 3).

**Figure 3:**
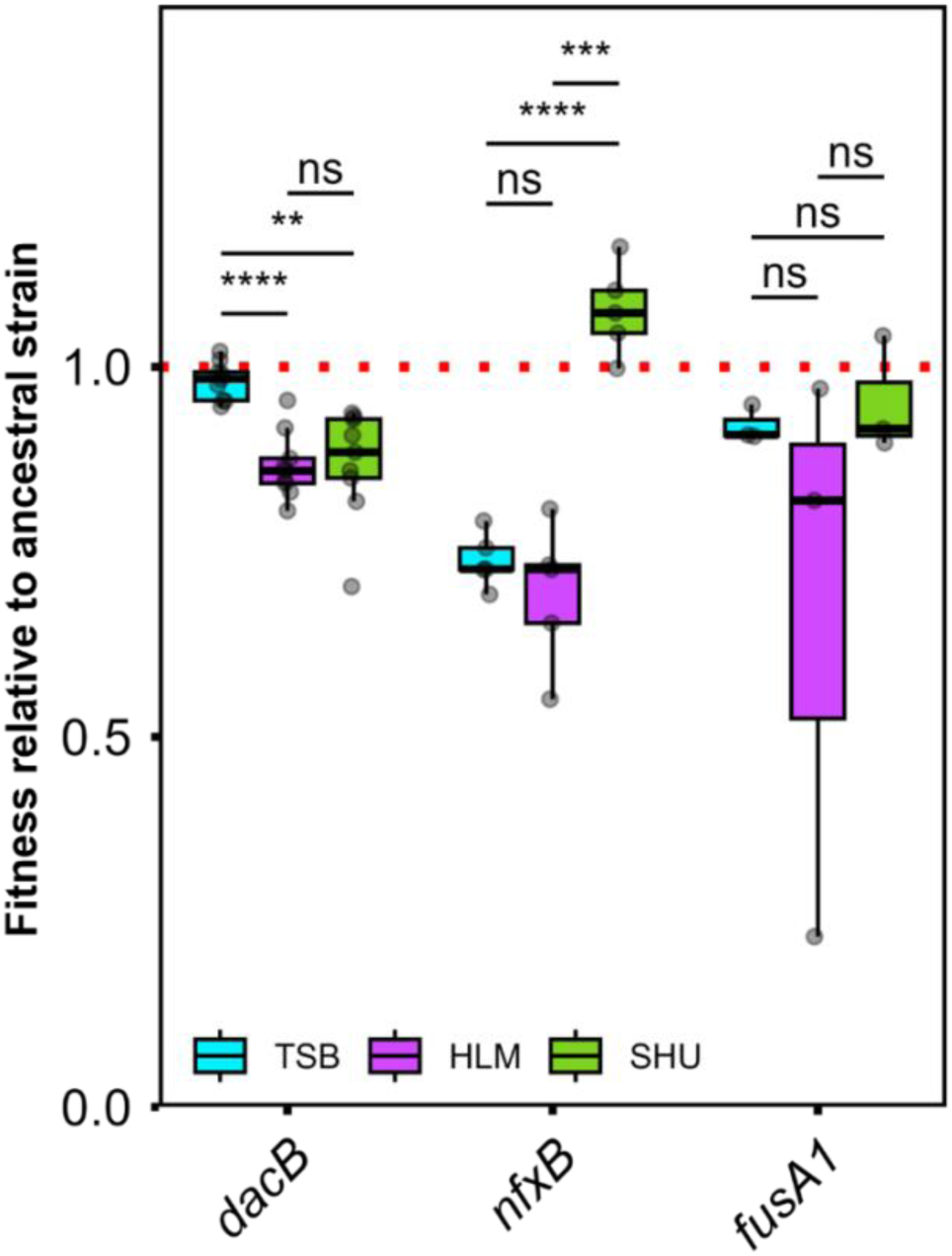
The impact of environment on the fitness costs associated with dacB, nfxB, and fusA1 resistance mutations. Fitness relative to the ancestral strain (ancestral fitness = 1.0, shown by red dotted line) was compared across environments for three groups: dacB mutants (PtzR1, PtzR2, PtzR3, PtzR6, CefR1, CefR3, CefR4, CefR5, CefR6), nfxB mutants (CipR1, CipR3, CipR4, CipR5, CipR6) and fusA1 mutants (AmiR1, AmiR3, AmiR5). Strains with mutations in more than one gene were excluded. Welch’s ANOVA followed by a Games–Howell post-hoc test was used to compare relative fitness across environments. Significance thresholds: p < 0.05 (*), p < 0.01 (**), p < 0.001 (***), p < 0.0001 (****).

The fitness costs associated with mutations in *dacB* and *nfxB* were significantly different in infection-like environments compared to in rich media (Figure 3). The relative fitness of *dacB-* mutants was significantly reduced in both HLM and SHU (Figure 3). The opposite effect was observed for mutations in *nfxB*, whereby *nfxB* mutants has significantly alleviated fitness costs in SHU, and the relative fitness of *nfxB-*mutants exceeded that of the ancestral strain (Figure 3). There were no significant differences in the fitness costs of *fus*A1 mutations across environments (Figure 3), although fitness costs incurred by the three *fusA1* mutations were highly variable in HLM (Figures 3, S3). We also conducted the same analysis grouping by selection antibiotic rather than single mutated gene (Figure S4).

### Mutations in *nfxB* provide a fitness benefit in the presence of oxalate

The *nfxB* mutants (cipR1-R6) had improved fitness in SHU, both relative to the ancestral strain in SHU and relative to their own growth in TSB (Figures 2B, 3 & S3). We screened components of SHU (oxalate, citrate, lactate, creatinine, urea and uric acid) to test for components that had different effects on the *nfxB* mutants compared to the ancestral strain (Figure 4), screening three independent *nfxB* mutants (cipR1, cipR2, cipR4) with different non-synonymous mutations (cipR1 and cipR4) and a deletion (cipR2).

**Figure 4:**
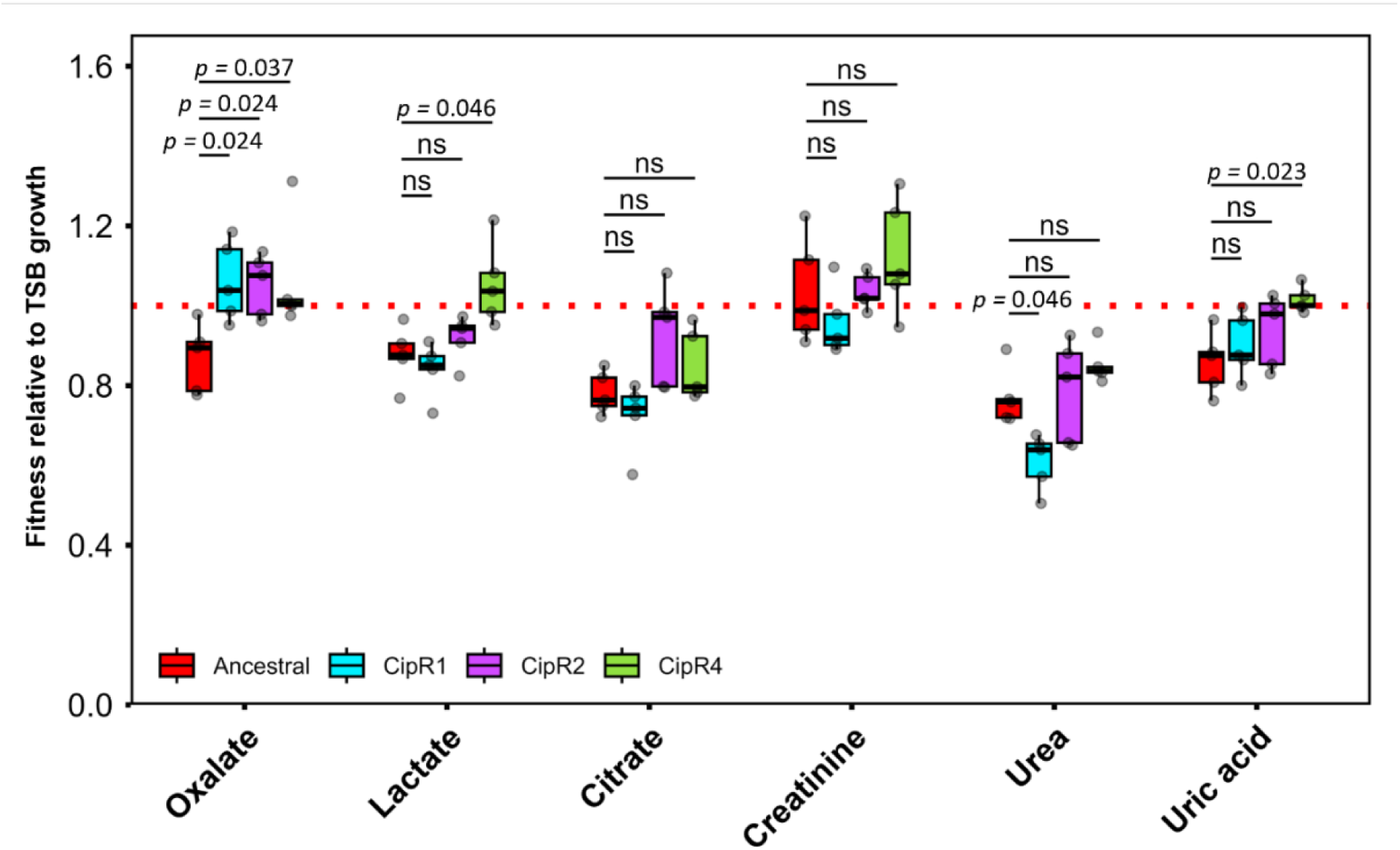
The impact of synthetic human urine components on nfxB mutant fitness compared to the ancestral strain. Relative fitness of each strain was calculated as the AUC in supplemented TSB divided by the AUC of the same strain and replicate in unsupplemented TSB (shown by the red dotted line), with 5 biological replicates per strain. The relative fitness of each mutant was compared to the ancestral strain. Due to heteroscedasticity and non-normal distribution of residuals, comparisons were analysed using a Welch’s ANOVA followed by post-hoc Welch’s t-tests with Bonferroni correction. Significance of p ≥ 0.05 = ns, otherwise p-value is shown.

The presence of oxalate improved the growth of all three *nfxB* mutants comparative to the ancestral strain (Figure 4). None of the other five components tested consistently improved *nfxB* mutant growth comparative to the ancestral strain (Figure 4). We also measured the impact of adjusting the pH of TSB to match that of SHU (5.6 pH) but found no significant differences in fitness costs between the strains related to pH (Figure S5).

## Discussion

A foundational concept in evolutionary biology is that the environment shapes the fitness of an organism. While the fitness costs of antibiotic resistance mutations represent a key barrier to the spread of resistance at an epidemiological scale, these costs have been largely quantified using *in vitro* measurements of bacteria in rich laboratory broth, such as LB or TSB^5,6^. Here, we demonstrate that the fitness costs of clinically important resistance mutations in *P. aeruginosa* change in both prevalence and magnitude in infection-like environments. The fitness costs of resistance mutations in rich media did not reliably predict those in infection-like environments (Figure 1, Figure 2B), and costs were on average increased in the lung-mimicking environment (Figure 2A), and less prevalent in the urinary tract-mimicking environment (Figure 2B). These findings are significant in relation to predicting the spread of antibiotic resistance, and highlight the importance of the chemical composition of infection- niches in shaping the fitness dynamics of resistance in *P. aeruginosa*.

Of the three possible growth environment combinations (TSB and HLM, TSB and SHU, HLM and SHU) we identified only one correlation in growth, which was between TSB and HLM (Figure 1). Although we identified consistency in the rank-order of growth and fitness in these environments, we also found that costly resistance mutations incurred a much greater magnitude of cost in HLM than in TSB (Figure 2), agreeing with meta-analysis suggesting the magnitude of fitness costs is higher when measured *in vivo* as opposed to *in vitro*^6^. When comparing costs in TSB to SHU, we identified no rank-order correlation and, in contrast to costs in HLM, costs in SHU were generally of a much lower magnitude, with the majority of the mutations found to incur no significant cost in this environment (Figures 2 & 3). Interestingly, our data showed that when the ancestral strain’s growth increased, the magnitude and prevalence of fitness costs also increased (Figures 2B & S3). This suggests that where the ancestral strain is stronger, there is a greater chance of a resistance mutation having a deleterious effect. This result supports predictions of Fisher’s geometric model, that fitness contours will curve tightly near the peak of the fitness landscape giving high costs of mutations in good environments^55^..

Within our panel of 33 resistant mutants, a subset of 17 mutants had resistance conferred by a mutation in only one of three genes (Table 1). These genes were *nfxB* (5 mutants), encoding the NfxB transcriptional repressor of the MexCD-OprJ multidrug efflux pump^43,44^; *dacB* (9 mutants), encoding a peptidoglycan-processing enzyme, in which mutations are characterised to lead to overexpression of the Amp-C beta-lactamase^41,42^; and *fusA1* (3 mutants), encoding elongation factor EF-G1A, in which mutations have been characterised as a mechanism of aminoglycoside resistance^45,46^. Mutations in *nfxB* were generated via selection on ciprofloxacin, mutations in *dacB* were generated via selection on either ceftazidime or piperacillin/tazobactam, and mutations in *fusA1* were generated via selection on amikacin (Table 1). The use of infection-like environments significantly altered the fitness costs of mutations in both *dacB* and *nfxB* (Figure 3). The *dacB* mutants exhibited greater costs in HLM and SHU than in TSB (Figure 3), and the *nfxB* mutants, which exhibited moderate fitness costs in TSB and HLM, exceeded the fitness of the ancestral strain in SHU (Figures 2B & 3). This suggests that *nfxB* mutants could have a selective advantage in the urinary tract environment and be selected for even in the absence of antibiotic treatment. This finding is particularly clinically significant; *nfxB* mutations are common in clinical *Pseudomonas* isolates, and ciprofloxacin is frequently used in the treatment of suspected *P. aeruginosa* infections^56^.

The MexCD-OprJ efflux pump can act on a wide variety of substrates in addition to clinical antibiotics^57–60^. Given that *nfxB* mutations confer resistance through the upregulation of this pump^44,51^, we hypothesised that the increase in relative fitness of *nfxB* mutants could be due to reduced sensitivity to components in SHU that negatively affect the growth of the ancestral strain, possibly driven by efflux via MexCD-OprJ. To test this hypothesis, we supplemented TSB with different SHU components that we thought could most plausibly have a negative impact on ancestral strain growth. These were nitrogenous waste products (creatinine, urea, and uric acid), organic acids (which would be present in carboxylate anion form in SHU; oxalate, lactate, and citrate), and reduced pH ^60–65^. The addition of all of these components, apart from creatinine and reduced pH, resulted in a notable reduction in ancestral strain growth (Figures 4 & S4). We identified one component, oxalate (supplemented via oxalic acid addition), in the presence of which *nfxB* mutations consistently provided significant benefits (Figure 4). Oxalic acid is an endogenously produced organic acid present in human urine in oxalate form at concentrations mimicked by SHU (∼16 mg/L)^15,66^, and is also a widespread organic acid across the biosphere^67^. Our findings suggest that the presence of oxalate in urine may impose a selective advantage on *P. aeruginosa* strains carrying *nfxB* mutations. This finding not only highlights a potential risk factor that may promote the spread of resistance during urinary tract infections, but also suggests an environmental variable that could possibly be manipulated to shape treatment outcomes.

In summary, using a panel of 33 unique *P. aeruginosa* resistant mutants generated against six clinically important antipseudomonal drugs, we identified clear differences between fitness costs in a standard laboratory medium (TSB) and infection-like environments (HLM, SHU), both in terms of magnitude and prevalence, and only one instance of correlation. The fitness costs of resistance mutations in rich media did not reliably predict those in infection-like environments (Figure 1, Figure 2B), and costs were on average increased in HLM and less prevalent in SHU (Figures 2A & 2B). These differences likely arise from the resistance mechanisms involved, including how they interact with local conditions and the extent to which costs can be offset by that environment. Furthermore, we have presented a specific example of how this occurs, with oxalate enhancing the fitness of *nfxB* mutants in SHU. Our results highlight the importance of environmental chemical composition for shaping the dynamics of antibiotic resistance in *P. aeruginosa* and supports the use of IMM as a simple way to the increase the physiological relevance of fitness cost assays.

## Author contributions

JK, LMP, CW, DRG, RCM, SJSC, and RMW contributed to data acquisition and data analysis. JK conducted all growth assays and fitness cost calculations. JK, LMP, LJK and RMW contributed to experimental design and methodology. SL, DRG, RCM, and RMW were responsible for generation and genome analysis of resistant mutants. JK and RMW were responsible for writing the original draft of the manuscript. DRG, RCM, SJSC, and LJK contributed to manuscript reviewing and editing. All authors read and approved the final manuscript.

## Acknowledgements

JK and LMP were supported by PhD studentships funded by the Department for the Economy (Northern Ireland). LJK has received funding from the European Union’s Horizon Europe research and innovation programme under the Marie Skłodowska-Curie grant agreement 101069111. RMW was supported by a UK Research and Innovation Future Leaders Fellowship (UKRI2317) and the Calleva Research Centre for Evolution and Human Sciences at Magdalen College (University of Oxford). We would like to thank MicrobesNG (Birmingham, UK) for the generation and initial processing of sequencing data. We would also like thank Dr Daniel Neill for his advice for preparing the HLM and mucin aliquot.

## Supplementary Information

**Table S1: Table of components used for each of the three growth environments (TSB, HLM and SHU).** Table includes the components listed in their final concentration. Tryptic soy broth (TSB) components were listed based on the manufacturers information. All components marked with an asterisk (*) were, in synthetic human urine (SHU), included as part of the pre-prepared SC Amino Acid mix and concentration are as detailed in the manufacturer’s information.

**Figure S1: Plate layout for growth measurements of *P. aeruginosa* in TSB, HLM, and SHU.** Shown are the plate layouts used for measuring mutants AztR4-CefR6. The same layout was used for other strains, in three further batches, comprised of AmiR1-AztR3, CefR1-CipR3, CipR4-PtzR6.

**Figure S2: Identification of resistant mutants’ fitness costs in each environment.** Comparisons between ancestral growth (AUC) on a per environment, to mutant growth, as summarised in Figure 2B. Datapoints for each replicate are shown by points, and mean is shown with a blue cross. The ancestral mean is represented by dotted red line. Here statistical significance was assessed using Welch’s ANOVA followed by a Games–Howell post hoc test, filtering only for ancestor to mutant comparisons. Significance annotations: p < 0.05 (*), p < 0.01 (**), p < 0.001 (***), p < 0.0001 (****).

**Figure S3: Changes in the fitness of strains across TSB, HLM, and SHU.** A) Comparison of the changes in relative fitness of resistant mutants in each environment, as assessed by their fitness relative to the ancestral strain in each environmental pair (TSB vs HLM, HLM vs SHU, SHU vs TSB), which is shown as a red dotted line. B) Comparison of changes in the growth (AUC) of strains, including only the ancestral strain (red dotted line) and those that had a significant change in relative fitness in Figure S2A. Statistical significance was assessed using Welch’s ANOVA followed by a Games–Howell post hoc test. Significance annotations: p < 0.05 (*), p < 0.01 (**), p < 0.001 (***), p < 0.0001 (****).

**Figure S2: The impact of environment on the fitness costs associated with the antibiotic used to select for mutant.** Fitness relative to the ancestral strain (shown by red dotted line at 1.0), was compared across environments, categorising mutants based on the selection antibiotic under which they mutated. Welch’s ANOVA followed by a Games–Howell post hoc test was used to compare relative fitness across environments. Significance annotations: p < 0.05 (*), p < 0.01 (**), p < 0.001 (***), p < 0.0001 (****).

**Figure S3: The impact of synthetic human urine pH on nfxB mutant fitness compared to the ancestral strain.** Relative fitness of each strain was calculated as the AUC in TSB adjusted to 5.6 pH divided by the AUC of the same strain and replicate in unsupplemented TSB (shown as a red dotted line), with 5 biological replicates per strain. The relative fitness of each mutant was compared to the ancestral strain. Comparisons were analysed using a Welch’s ANOVA followed by post hoc Welch’s t-tests with Bonferroni correction. Not significant (ns) = p ≥ 0.05.

## Supplementary Data

**Supplementary Data 1:** AUC data for strains in TSB, SHU, and HLM.

**Supplementary Data 2:** SHU component supplementation AUC data.

