## Supplementary Information for "Fitness costs of antibiotic resistance mutations in rich media do not reliably predict those in infection-like environments"

***Table S1: Table of components used for each of the three growth environments (TSB, HLM and SHU).*** *Table includes the components listed in their final concentration. Tryptic soy broth (TSB) components were listed based on the manufacturers information. All components marked with an asterisk (*) were, in synthetic human urine (SHU), included as part of the pre-prepared SC Amino Acid mix and concentration are as detailed in the manufacturer's information.*

| Description | Component | TSB | HLM | SHU |
|---|---|---|---|---|
| **Carbohydrate metabolites** | Galactose | - | 6.27 μg/L | - |
| | Glucose | 2.5 g/L | 4 g/L | - |
| | Myo-Inositol* | - | - | 85.6 mg/L |
| | N-acetyl glucosamine | - | 1.28 g/L | - |
| | Sialic acid | - | 3.23 μg/L | - |
| **Nucleic acids/bases** | eDNA | - | 0.96 g/L | - |
| | Adenine* | - | - | 21 mg/L |
| | Uracil* | - | - | 85.6 mg/L |
| **Host-derived antimicrobials** | Lactoferrin | - | 1 mg/L | - |
| | Lysozyme | - | 8 mg/L | - |
| **Host-derived macromolecules** | Albumin | - | 1.5 g/L | - |
| | Mucin | - | 1.2 g/L | - |
| **Peptones** | Casein peptone | 17 g/L | - | - |
| | Soya peptone | 3 g/L | - | - |
| **Amino acids** | L-Alanine* | - | 3 mg/L | 86.5 mg/L |
| | L-Arginine | - | 37 mg/L | - |
| | L-Arginine HCl* | - | - | 85.6 mg/L |
| | L-Asparagine* | - | 0.35 mg/L | 85.6 mg/L |
| | L-Aspartic acid* | - | 1 mg/L | 85.6 mg/L |
| | L-Cysteine | - | 0.28 mg/L | - |
| | L-Cysteine HCl* | - | - | 85.6 mg/L |
| | L-Glutamine* | - | 0.5 mg/L | 85.6 mg/L |
| | L-Glutamic acid* | - | - | 85.6 mg/L |
| | Glycine* | - | 2.7 mg/L | 85.6 mg/L |
| | L-Histidine | - | 0.5 mg/L | - |
| | L-Histidine HCl* | | - | 85.6 mg/L |
| | L-Isoleucine* | - | 1 mg/L | 85.6 mg/L |
| | L-Leucine* | - | 2mg/L | 173.4 mg/L |
| | L-Lysine | - | 2.2 mg/L | - |
| | L-Lysine HCl* | - | - | 85.6 mg/L |
| | L-Methionine* | - | 0.3 mg/L | 85.6 mg/L |
| | L-Phenylalanine* | - | 0.8 mg/L | 85.6 mg/L |
| | L-Proline* | - | 1.3 mg/L | 85.6 mg/L |
| | L-Serine* | - | 1.8 mg/L | 85.6 mg/L |
| | L-Threonine* | - | 15 mg/L | 85.6 mg/L |
| | L-Tryptophan* | - | 0.16 mg/L | 85.6 mg/L |
| | L-Tyrosine* | - | 0.73 mg/L | 85.6 mg/L |
| | L-Valine* | - | 1.6 mg/L | 85.6 mg/L |
| **Nitrogenous waste products** | Creatinine | - | - | 1.018 g/L |
| | Uric acid | - | - | 0.101 g/L |
| | Urea | - | - | 16.817 g/L |

| | | | | |
|---|---|---|---|---|
| **Polyamines** | Putrescine | - | 616 µg/L | - |
| | Spermidine | - | 0.25 mg/L | - |
| | Spermine | - | 44.5 µg/L | - |
| **Organic acids / acid salts** | Oxalic acid | - | - | 16 mg/L |
| | Para-aminobenzoic acid* | | - | 8.6 mg/L |
| | Succinate | - | 295 µg/L | - |
| | Sodium L-lactate | - | - | 0.123 g/L |
| | Trisodium citrate dihydrate | - | - | 2.264 g/L |
| **Salts** | Ammonium chloride | - | - | 1.07 g/L |
| | Calcium chloride | - | 45 mg/L | 0.444 g/L |
| | Copper(II) chloride | - | 106 µg/L | - |
| | Iron(II) chloride | - | 887 µg/L | - |
| | Iron(II) sulfate | - | - | 1 mg/L |
| | Magnesium chloride | - | 8 mg/L | - |
| | Magnesium chloride hexahydrate | - | - | 0.651 g/L |
| | Magnesium sulfate | - | - | 0.385 g/L |
| | Potassium chloride | - | - | 2.833 g/L |
| | Potassium phosphate monobasic | - | - | 2.177 g/L |
| | Disodium hydrogen phosphate dihydrate | - | - | 1.157 g/L |
| | Dipotassium hydrogen phosphate | 2.5 g/L | - | - |
| | Sodium phosphate monobasic | - | - | 0.648 g/L |
| | Sodium bicarbonate | - | - | 1.134 g/L |
| | Sodium chloride | 5 g/L | 1 g/L | 5.844 g/L |
| | Sodium sulfate | - | - | 2.415 g/L |
| | Zinc chloride | - | 390 µg/L | - |
| **Acid/Base for pH adjustment** | Potassium hydroxide | - | As | - |
| | Hydrochloric acid | - | - | As needed |
| | Sodium hydroxide | - | - | As needed |
| | **Adjusted pH:** | **7.3 ±0.2** | **6.85** | **5.6** |

| | 1 | 2 | 3 | 4 | 5 | 6 | 7 | 8 | 9 | 10 | 11 | 12 |
|---|---|---|---|---|---|---|---|---|---|---|---|---|
| A | | | | | | | | | | | | |
| B | | AztR4 | AztR6 | CefR2 | CefR4 | CefR6 | **Ancestor** | CefR5 | CefR3 | CefR1 | AztR5 | |
| C | | AztR4 | AztR6 | CefR2 | CefR4 | CefR6 | **Ancestor** | CefR5 | CefR3 | CefR1 | AztR5 | |
| D | | AztR4 | AztR6 | CefR2 | CefR4 | CefR6 | **BLANK** | CefR5 | CefR3 | CefR1 | AztR5 | |
| E | | AztR5 | CefR1 | CefR3 | CefR5 | **BLANK** | CefR6 | CefR4 | CefR2 | AztR6 | AztR4 | |
| F | | AztR5 | CefR1 | CefR3 | CefR5 | **Ancestor** | CefR6 | CefR4 | CefR2 | AztR6 | AztR4 | |
| G | | AztR5 | CefR1 | CefR3 | CefR5 | **Ancestor** | CefR6 | CefR4 | CefR2 | AztR6 | AztR4 | |
| H | | | | | | | | | | | | |

| | 1 | 2 | 3 | 4 | 5 | 6 | 7 | 8 | 9 | 10 | 11 | 12 |
|---|---|---|---|---|---|---|---|---|---|---|---|---|
| A | | | | | | | | | | | | |
| B | | AztR4 | AztR6 | CefR2 | CefR4 | CefR6 | **Ancestor** | CefR5 | CefR3 | CefR1 | AztR5 | |
| C | | AztR4 | AztR6 | CefR2 | CefR4 | CefR6 | **Ancestor** | CefR5 | CefR3 | CefR1 | AztR5 | |
| D | | AztR4 | AztR6 | CefR2 | CefR4 | CefR6 | **BLANK** | CefR5 | CefR3 | CefR1 | AztR5 | |
| E | | AztR5 | CefR1 | CefR3 | CefR5 | **BLANK** | CefR6 | CefR4 | CefR2 | AztR6 | AztR4 | |
| F | | AztR5 | CefR1 | CefR3 | CefR5 | **Ancestor** | CefR6 | CefR4 | CefR2 | AztR6 | AztR4 | |
| G | | AztR5 | CefR1 | CefR3 | CefR5 | **Ancestor** | CefR6 | CefR4 | CefR2 | AztR6 | AztR4 | |
| H | | | | | | | | | | | | |

| | 1 | 2 | 3 | 4 | 5 | 6 | 7 | 8 | 9 | 10 | 11 | 12 |
|---|---|---|---|---|---|---|---|---|---|---|---|---|
| A | | | | | | | | | | | | |
| B | | AztR4 | AztR6 | CefR2 | CefR4 | CefR6 | **Ancestor** | CefR5 | CefR3 | CefR1 | AztR5 | |
| C | | AztR4 | AztR6 | CefR2 | CefR4 | CefR6 | **Ancestor** | CefR5 | CefR3 | CefR1 | AztR5 | |
| D | | AztR4 | AztR6 | CefR2 | CefR4 | CefR6 | **BLANK** | CefR5 | CefR3 | CefR1 | AztR5 | |
| E | | AztR5 | CefR1 | CefR3 | CefR5 | **BLANK** | CefR6 | CefR4 | CefR2 | AztR6 | AztR4 | |
| F | | AztR5 | CefR1 | CefR3 | CefR5 | **Ancestor** | CefR6 | CefR4 | CefR2 | AztR6 | AztR4 | |
| G | | AztR5 | CefR1 | CefR3 | CefR5 | **Ancestor** | CefR6 | CefR4 | CefR2 | AztR6 | AztR4 | |
| H | | | | | | | | | | | | |

TSB    HLM    SHU

***Figure S1: Plate layout for growth measurements of P. aeruginosa in TSB, HLM, and SHU.*** *Shown are the plate layouts used for measuring mutants AztR4-CefR6. The same layout was used for other strains, in three further batches, comprised of AmiR1-AztR3, CefR1-CipR3, CipR4-PtzR6.*

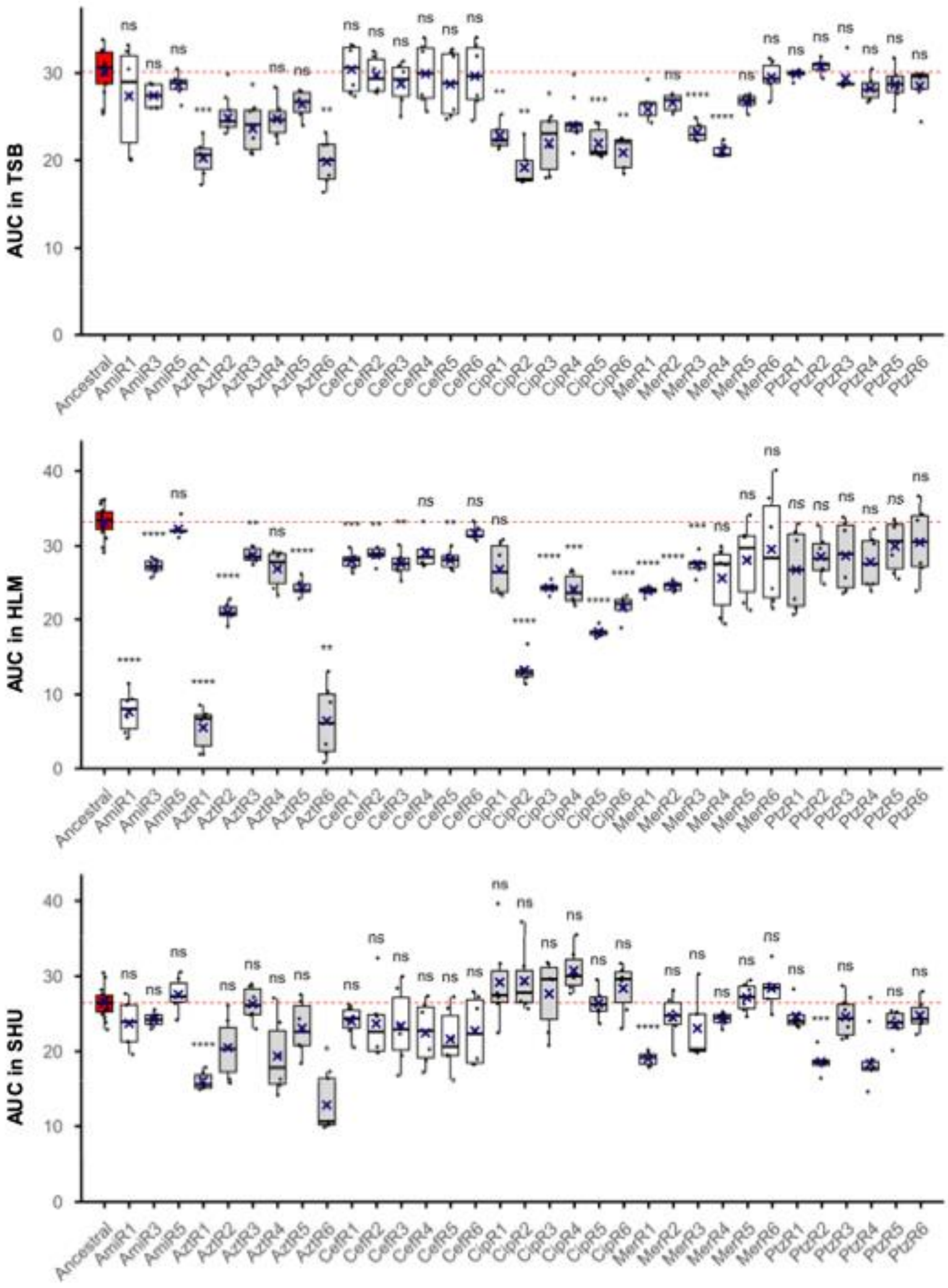


***Figure S2: Identification of resistant mutants' fitness costs in each environment.*** *Comparisons between ancestral growth (AUC) on a per environment, to mutant growth, as summarised in Figure 2B. Datapoints for each replicate are shown by points, and mean is shown with a blue cross. The ancestral mean is represented by dotted red line.* Here s*tatistical significance was assessed using Welch's ANOVA followed by a Games–Howell post hoc test, filtering only for ancestor to mutant comparisons. Significance annotations: $p < 0.05$ (*), $p < 0.01$ (**), $p < 0.001$ (***), $p < 0.0001$ (****).*

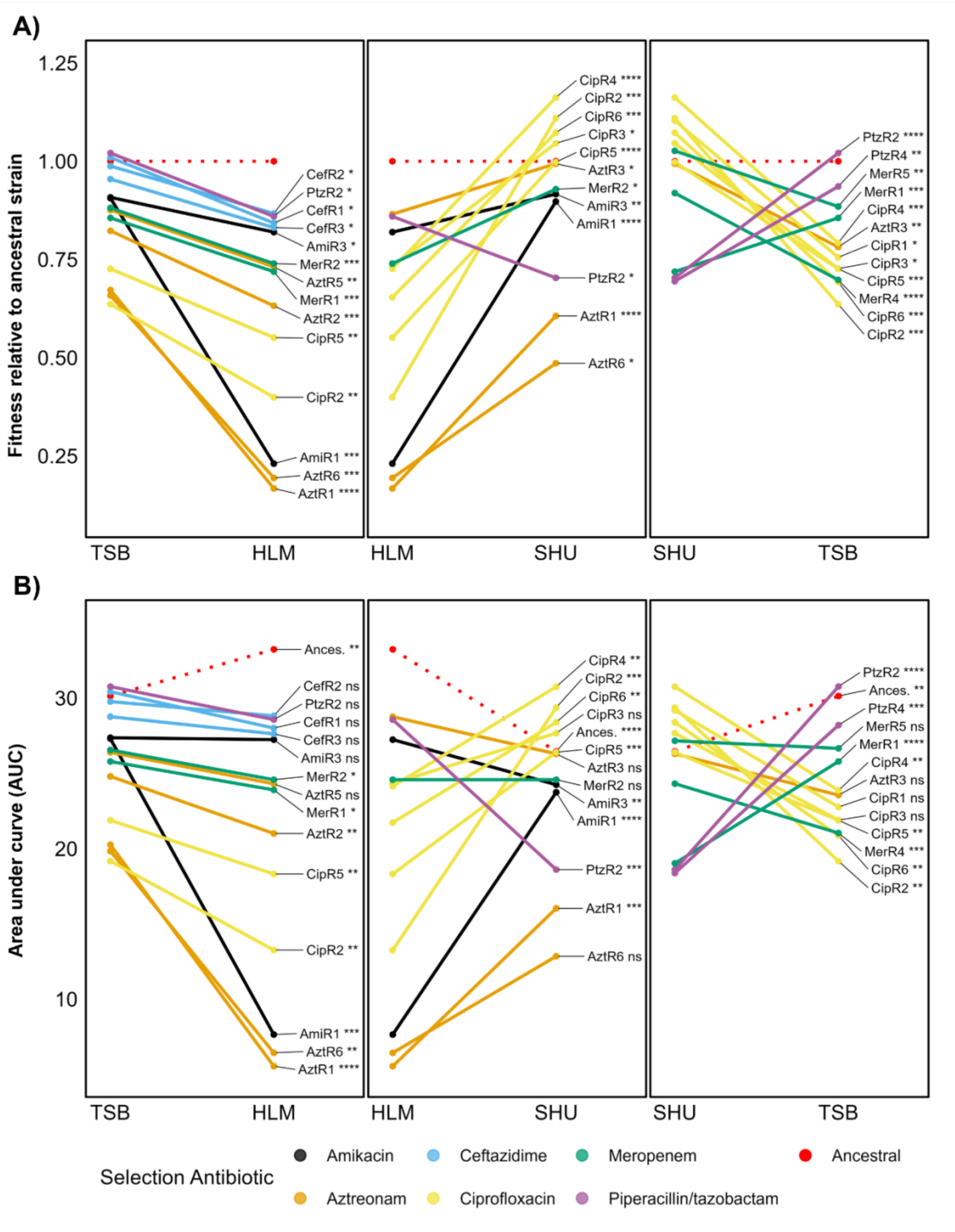


***Figure S3: Changes in the fitness of strains across TSB, HLM, and SHU.*** *A) Comparison of the changes in relative fitness of resistant mutants in each environment, as assessed by their fitness relative to the ancestral strain in each environmental pair (TSB vs HLM, HLM vs SHU, SHU vs TSB) , which is shown as a red dotted line. B) Comparison of changes in the growth (AUC) of strains, including only the ancestral strain (red dotted line) and those that had a significant change in relative fitness in Figure S2A. Statistical significance was assessed using Welch's ANOVA followed by a Games–Howell post hoc test. Significance annotations: p < 0.05 (*), p < 0.01 (**), p < 0.001 (***), p < 0.0001 (****).*

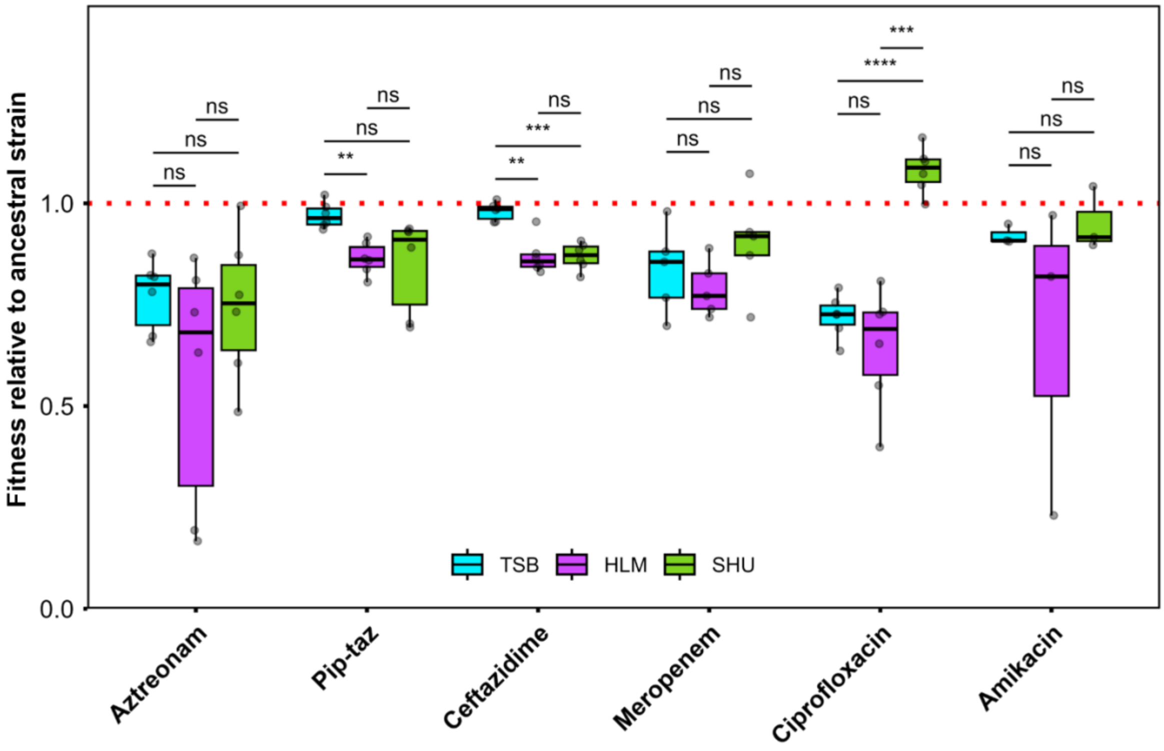


***Figure S2: The impact of environment on the fitness costs associated with the antibiotic used to select for mutant.*** *Fitness relative to the ancestral strain (shown by red dotted line at 1.0), was compared across environments, categorising mutants based on the selection antibiotic under which they mutated. Welch's ANOVA followed by a Games–Howell post hoc test was used to compare relative fitness across environments. Significance annotations: p < 0.05 (*), p < 0.01 (**), p < 0.001 (***), p < 0.0001 (****).*

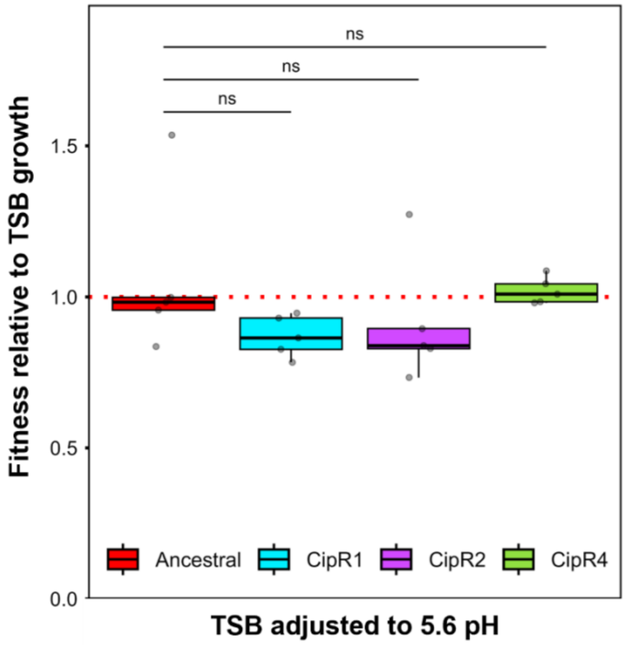


***Figure S3: The impact of synthetic human urine pH on nfxB mutant fitness compared to the ancestral strain.*** *Relative fitness of each strain was calculated as the AUC in TSB adjusted to 5.6 pH divided by the AUC of the same strain and replicate in unsupplemented TSB (shown as a red dotted line), with 5 biological replicates per strain. The relative fitness of each mutant was compared to the ancestral strain. Comparisons were analysed using a Welch's ANOVA followed by post hoc Welch's t-tests with Bonferroni correction. Not significant (ns) = p ≥ 0.05.*
